## Supplementary Information for "Short- and long-read metagenomics of urban and rural South African gut microbiomes reveal a transitional composition and novel taxa"

#### Community engagement

It is an ethical imperative to include diverse and understudied populations in biomedical research, and it is of utmost importance to conduct research equitably in groups which may be socioeconomically vulnerable. During the course of this project, researchers traveled to field sites in both Soweto and Bushbuckridge on multiple occasions, including a trip to the Agincourt Health and Demographic Surveillance Site (HDSS) early in the planning phases of the study where researchers met with members of the Community Advisory Group (CAG). This team of community members from across the Bushbuckridge Municipality routinely meets with researchers to discuss and approve research projects in the community and voice questions and concerns on behalf of the community. The research team sought the opinion of the CAG as to whether the study of stool would be appropriate in the community, given that attitudes toward collection of fecal samples may differ across cultures. The CAG expressed that it would be appropriate to collect stool if it was explained clearly to participants that the stool was for scientific purposes, that it would not be used to cause harm in any way to participants, and that it would not be sold.

On a second visit to the Agincourt HDSS, the research team conducted a microbiome workshop for community members to attend to learn more about how studying the gut microbiome may improve understanding of the health of African populations. Community members learned about bacteria, their role in human health, and the ways in which studying gut microbiome composition might impact our understanding of human health in the region. Community members were given the opportunity to ask questions and discuss the research project.

### Additional information on study sites

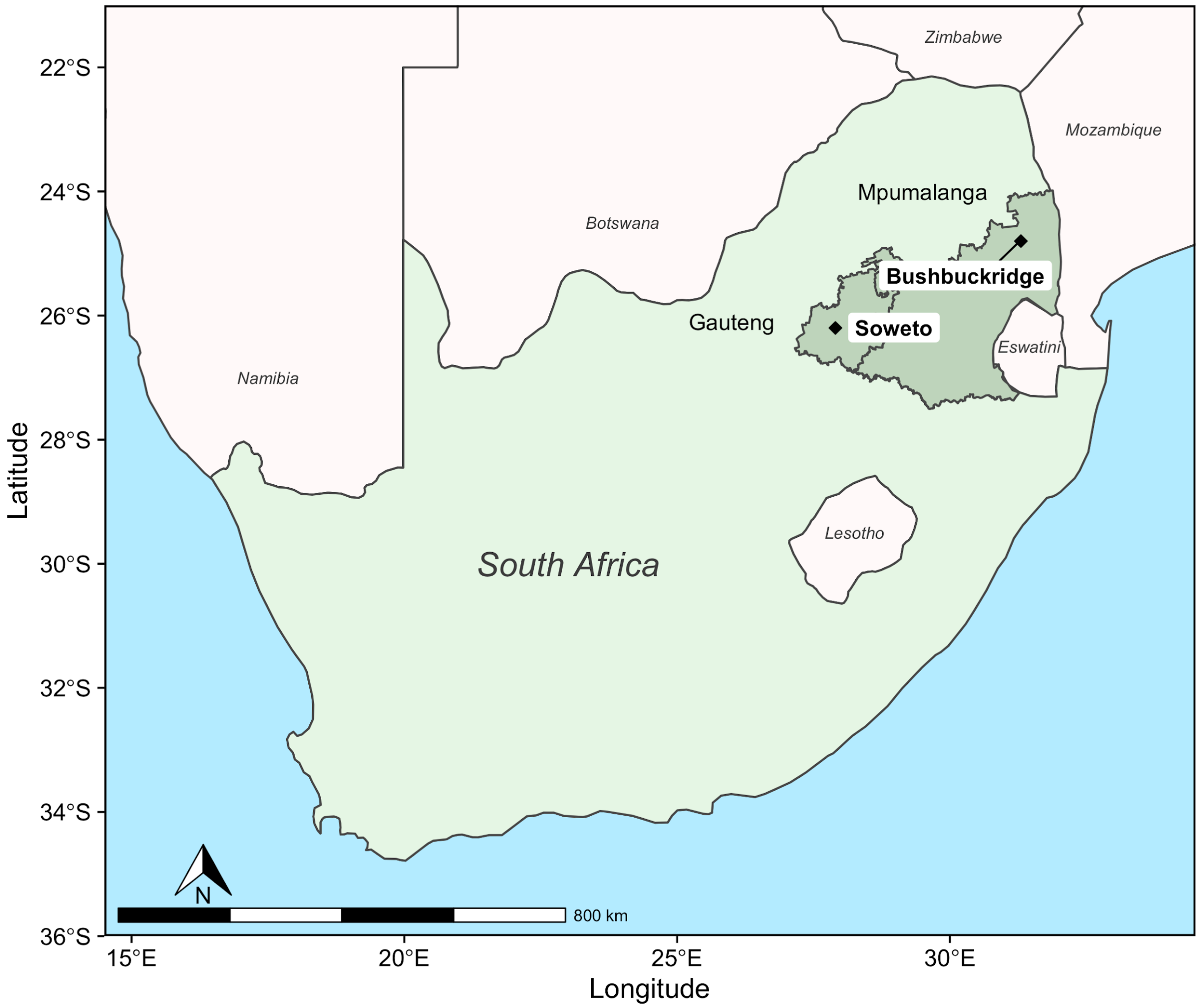


South Africa is a prime example of a country undergoing rapid lifestyle and epidemiological transition. With the exception of the HIV/AIDS epidemic in the mid-1990s to the mid-2000s, over the past three decades South Africa has experienced a steadily decreasing mortality from infectious disease and an increase in noncommunicable disease[1,2]. Concomitantly, increasingly sedentary lifestyles and changes in dietary habits, including access to calorie-dense processed foods, contribute to a higher prevalence of obesity in many regions of South Africa[2], a trend which disproportionately affects women[3,4].

The University of the Witwatersrand in Johannesburg, South Africa has long-standing partnerships with health and demographic surveillance sites (HDSSs) across

South Africa. The Agincourt HDSS, run by the MRC/Wits Rural Public Health and Health Transitions Research Unit surveys individuals living in the Bushbuckridge Municipality in rural Mpumalanga province. Bushbuckridge Municipality is the largest municipality in Mpumalanga by population and by land area (10,250 $km^2$), and includes areas of Kruger National Park. The MRC/Wits Developmental Pathways for Health Research Unit surveys individuals living in urban Soweto, a township of 200.03 $km^2$ that is part of Johannesburg in Gauteng province. Soweto's name is derived from **So**uth **We**stern **To**wnships and is formally incorporated in the city of Johannesburg. The populations of both Bushbuckridge and Soweto are predominantly black African (http://www.statssa.gov.za)

As a result of HDSS surveillance, much is known about changes in disease risk and causes of death over time in Bushbuckridge and Soweto. For instance, rates of obesity are higher in women compared to men in both Bushbuckridge and Soweto HDSS surveillance areas[5,6].

Additionally, diet and lifestyle in Bushbuckridge and Soweto have been extensively surveyed. Pisa et al.[7] administered food frequency questionnaires to adolescents living within the Agincourt HDSS (Bushbuckridge) and identified nutrient patterns via principal components analysis that explain 79% of the variance in nutritional intake in the study population. They found that female gender and being in the lowest socioeconomic status (SES) tertile were associated with animal driven nutrients, being in mid-puberty associated with vitamins, fiber, and vegetable oil nutrients, and physical activity and being in the lowest SES tertile were associated with mixed diet driven nutrients. Additionally, body mass index (BMI) was found to associate with nutritional patterns. Sedibe et al.[8] surveyed dietary practices and rates of obesity in adolescents from Bushbuckridge and Soweto and found that across study sites, participants regularly consumed fast foods. Rural and/or male participants were less likely to be overweight and obese than females, and irregular consumption of breakfast on weekdays was associated with increased risk of overweight and obesity. A qualitative study of adolescent females in Bushbuckridge described dietary practices. Findings included the observation that pap (maize porridge) and tea are common breakfast options among the population of Bushbuckridge, and that leafy vegetables, legumes, and nuts are viewed as healthy by participants but affordability can be a barrier to access, and fruit is not easily accessible. Participants described that limitations in household resources restricted their options for purchasing healthy foods at school, and that instead they bought cheaper options including bread, vetkoek (fried bread), kota (white bread filled with potato chips and processed meat or cheese) or other processed snack foods. In adult South Africans participating in AWI-Gen, consumption of sugar-sweetened beverages was positively associated with BMI in women.[9]

#### Microbiome and human genetic association testing

To test whether Bray-Curtis distance was correlated with genetic distance, we used 114 samples from Bushbuckridge for whom QC had passed on both human and microbiome data. This is a low *n*, so the absence of correlation in this study is only weak evidence of a general claim of lack of correlation. We did not include the Soweto data because the different environment would have been a confounding factor. All participants in the AWI-Gen study have been genotyped using the H3A Custom Array, an approximately 2.2 million SNP array designed to maximise coverage in African populations (https://h3abionet.org/h3africa-chip). QC of the genotype data was done using the H3ABioNet H3AGWAS QC pipeline (https://github.com/h3abionet/h3agwas). Genetic relatedness (PI-HAT) was computed using PLINK [48] and converted to a distance score. We also computed principal components using PLINK. Data for our participants was extracted. There was no significant correlation between the PI-HAT distance matrix and the Bray-Curtis distance (Mantel test, $10^5$ permutations). As a supplementary test, using the same Mantel test, we computed correlation between the Bray-Curtis distance and the first ten principal components. The highest correlation was with PC3 with a correlation of 0.13 and $p$=0.027. However, not only is this correlation low, after correcting for multiple testing, the $p$ value is above any reasonable cut-off.

Association testing was done using the same human genetic data against abundance levels of *Prevotella copri*, *Escherichia coli*, *Alistipes sp* CAG-435, *Faecalibacterium prausnitzii*, *Bacteroides vulgatus*, *Prevotella sp* TF12-30, *Prevotella sp* AM23-5, *Prevotella sp* AM42-24, *Bacteroides fragilis*, *Ruminococcaceae bacterium*. Again, because of the environmental difference between Bushbuckridge and Soweto we only used the 114 Bushbuckridge samples who passed QC both on human and microbiome data. Due to the small data size, we filtered out all SNPs with minor allele frequency less than 5%. Imputation was done using the Sanger imputation service using the African Genome Resource reference panel, yielding 7.757 million well imputed SNPs. None of the abundance levels had normal distributions, and in some cases there were extreme abundance levels far from the mean, which could bias the results given our small sample size. For this reason we log-transformed the abundance levels. We used the H3Agwas Association Testing pipeline (github.com/h3abionet/h3agwas) using GEMMA[10] as the underlying association testing tool. We used $5x10^{-8}$ as the *p*-value to account for multiple testing both with multiple SNPs and microbiome genera. The QQ-plots for all genera tested were well-behaved except for *Ruminococcaceae bacterium*, which we excluded from further analysis. Eight SNPs were below the cut-off, with several more in the suggestive level. This is shown in Table S6 which shows the genus, SNP, location of the SNP, the β-value and the p-value. Since we have a small sample and the *p*-values are not very far below the cut-off we present the results here as an initial analysis that might be useful for follow-up work.

The most interesting hit is in the *SLC2A10* gene, which encodes for a glucose transporter. Besides the two SNPs listed in the table there were several other SNPs in the same gene which had *p*-values above the cut-off but in a suggestive region. SNP rs10137347 is in a transcription factor binding site; the gene it is most associated with is *RNASE6* gene, which encodes a protein that is active in the urinary tract and which has antimicrobial properties, particularly against gram-negative bacteria (which Alistipes is). *FOXP1* is a gene with complex functions, but recent work shows that it is implicated in regulating the immune system. Finally, the GFRA1 protein is a neurotrophic factor, which is difficult to associate with the microbiome functioning.
